# Flexible use of contact calls in a species with high fission-fusion dynamics

**DOI:** 10.1101/2022.05.25.492876

**Authors:** Briseño-Jaramillo Margarita, Sosa-López José Roberto, Ramos-Fernández Gabriel, Lemasson Alban

## Abstract

The ‘social complexity hypothesis’ posits that complex social systems (that entail high uncertainty) require complex communicative systems (with high vocal flexibility). In species with fission-fusion dynamics, where the fluid composition of temporary subgroups increases the uncertainty with which group members must manage their social relationships, vocal communication must be particularly flexible. This study assessed whether contact call rates vary with caller and audience characteristics in free-living spider monkeys, as well as with fission and fusion events. Adult females and immature individuals called more when in small audience settings, while audience size did not influence adult males. Adults called more when in the presence of the opposite sex, whereas immatures vocalized more in subgroups composed only by females. Females also called more when with their mature sons. We found higher call rates in periods during which fission and fusion events took place than in periods with more stable compositions and when the composition after a fission or fusion event changed from one sex to two sexes. A flexible use of contact calls allows individuals to identify themselves when they join others, particularly if they are members of the opposite sex. This socio-spatial cohesion function reduces the uncertainty about subgroup composition.

## INTRODUCTION

According to the “social complexity hypothesis for communicative complexity”, patterns of communication and patterns of social organization are functionally related [1,2]. Animals living in complex social groups should thus use complex communication systems. Several studies have shown support for this hypothesis in the case of vocal signals from a broad range of taxa, including primates, with an impact of social complexity on vocal repertoire composition [3–5], acoustic diversity and plasticity [6–8], vocal exchange rules [9] and context-dependent call rate flexibility [10,11]. However, given that the motor control that non-human primates have over their vocal tracts is limited [12] and that their vocal repertoires are composed by a limited number of call types [13,14], an effective way to deal with this problem, in the context of a complex social environment, is to use the same type of vocalization in a flexible way to transmit different messages or to fulfill different social functions. Evidence of vocal flexibility under social influences (see also “pragmatic flexibility” in [15]) has been provided in several nonhuman primate species [16,17].

While there is no consensus about the definition of social complexity [18], it is widely accepted that uncertainty is a prominent characteristic of complex systems [11]. Fission-fusion dynamics, a property of animal groups that split in temporary aggregations (subgroups from here on; [19], can lead to a high variability in subgroup composition. This variability implies a highly variable social context in which any group member may find itself. In an effort to quantify this variability in subgroup composition, Ramos-Fernandez et al. [20] used information theoretic measures of the amount of uncertainty found in the composition of subgroups formed by different species. This uncertainty is higher in species with high degrees of fission-fusion dynamics, like spider monkeys (*Ateles geoffroyi*) and chimpanzees (*Pan troglodytes*), than in multi-level species that form variable subgroups but with a more predictable composition, like geladas (*Theropitecus gelada*). In spider monkeys and chimpanzees, individuals experience a high level of uncertainty about the particular set of associates they have at any given time. Thus, it is expected that these species will have developed behavioral mechanisms to reduce this uncertainty [20,21].

The vocal repertoire of nonhuman primates is typically composed of several acoustically and functionally distinctive types of calls. Some of these call types have a clearly identifiable function, such as alarm calls (associated with the presence of danger) [22], threat or distress calls (associated with social conflicts) [23], or food calls (associated with foraging and feeding) [24,25]. Other call types, are emitted at any time of day in a wide variety of contexts [26]. Within this category is the so-called “contact call”, the signal type with the highest individual acoustic variability within the repertoire [27]. In most nonhuman primates, these call types may function for regulating spatial cohesion facilitating the location of group members by auditory signals, an essential requirement for species living in visually dense habitats [26,28] and that experience high uncertainty in the composition of their subgroups, as mentioned above. But these calls may also function, in a non-mutually exclusive way, in regulating social cohesion by facilitating and maintaining social bonds [29]. In fluid societies, identifying callers at a distance could play an important role in maintaining social relationships by reducing the uncertainty about the presence/absence of important partners.

There is evidence that shows that spider monkeys use contact calls for both spatial and social cohesion. It is known that one of these calls, the whinny call, contains information about caller identity and they have been shown to allow individuals to obtain information about subgroup members’ locations[30,31]. They can serve a spacing function by which callers regulate their positions relative to each other through an evaluation of potential consequences based on their relationships [32]. Whinnies could thus help to mediate interindividual spacing during contexts where too much proximity increases feeding competition, and this seems particularly true for females who experience a lot of competition and call more than males during foraging [32]. In line with that, there is evidence that contact call rates, more so again for females, are positively correlated with the number of individuals joining or leaving their subgroup [32]. In addition, playback experiments showed that whinnies can promote the approach of close associates of a caller compared to other, less closely related, individuals [30,31]. Whinnies are loud enough to be audible by any member of a given subgroup and beyond [30]. Ordóñez-Gómez et al. [33] showed that individuals even lower the fundamental frequency of whinnies to facilitate communication and to maintain contact with distant individuals (those traveling outside their subgroup). Social cohesion is also supported by Briseño-Jaramillo et al. [34] showing that call-matching between two individuals is higher between preferred grooming partners, in line with the Dunbar’s “grooming-at-distance” hypothesis [35].

The evidence points at the whinny’s main function being related to regulating subgroup composition, by promoting the cohesion between callers and their closely associated recipients, and at the same time the repulsion of other recipients of the same call who are not closely associated with the caller. It is not yet clear, however, how individuals use their calls in contexts that vary in the uncertainty levels (e.g. during fusion, fission or stable periods). During these changing periods, could the whinnies function in maintaining sex-segregation and regulating sex-encounters? During stable periods, could whinnies function in the coordination of interindividual distances to different audience compositions? If so, will it be reflected in call rates at the individual and subgroup level? And finally, it is not known whether the presence of a specific individual in the audience modifies individual vocal behavior.

In this study, we take advantage of the large variability in subgroup composition shown by the species to evaluate how vocal behavior at the individual and subgroup levels varies between different social contexts. We evaluate whether different age-sex classes flexibly use contact calls in diverse social contexts and whether fission and fusion events, when subgroup composition changes, are also associated with differences in calling behavior. Spider monkeys occupy large areas of dense forest habitats with patches of fruit trees that change seasonally [36]. They live in groups with a high degree of fission–fusion dynamics, where the size and spatial cohesion of subgroups is dynamically adjusted depending on the availability and distribution of food [37,38]. Sex is clearly a structuring factor of these subgroups, as its more common to observe single-than mixed-sex subgroups [39].

Adult males are philopatric and more socially active than adult females [40], developing strong, long-lasting relationships with one another and cooperating to compete with neighboring groups’ males [41]. Conversely, adult females disperse from their natal group and do not establish strong bonds with kin group members [42]. Thus, males are keen to maintain social bonds, while females are less gregarious [43]. Hence, the value of adult female relationships is assumed to be low [44], as they spend less time with other group members than adult males and form smaller subgroups or forage alone with dependent offspring [45,46], interacting affilatively less than males do amongst them [39]. Mixed-sex relationships involve high rates of aggression from males to females, including coalitionary behaviors between males when they are in mixed-sex subgroups [39]. However, male-female relationships can vary in different circumstances (e.g. the reproductive status of females) [43] or kinship (e.g. long-term associations between mothers and mature males offspring) [47].

Specifically, we studied how individuals of different age/sex classes flexibly modify their vocal behavior according to the social context by testing how contact call rates vary according to three factors: subgroup size and composition, and presence of specific individuals in the subgroup. We also studied how fission and fusion events affect the calling rate at the level the subgroup, by comparing their calling rate during periods when subgroups come together and they split in two, compared to other periods when no fission or fusion event occurs.

We hypothesized that 1) if contact calls are used to regulate inter-individual distances (notably via the attraction and repulsion of conspecifics, depending on their social relationships), then subgroup size should differentially affect male and female calling behavior. Because competition (notably for food) affects females more than males [48] and because they are less invested than males in the regulation of social relationships in general [44], we predicted that large subgroups would inhibit female calling but stimulate male calling. Due to strong maternal influence, we expected that immature individuals would present the same pattern as mature females. Second, whinnies could help in the regulation of subgroup composition and notably the presence of the same or opposite sex, as callers are approached by close associates and potentially avoided by others [32]. We hypothesized that, 2) if whinnies function in coordinating sex-segregation and regulating sex-encounters, contact call rates would depend on the sexual composition of the audience. We expected call rates in adult females and males to be affected by the presence of the opposite sex in the audience, especially in the case of males, who have been shown to engage in higher coalition rates with other males when in the presence of females. Due to the lack of sexual maturity, we also did not expect immatures to be affected by the presence of the opposite sex in the same way than mature sex subgroups, compared to other times when fusions led to the same sex-based composition or when there were no fusion events.

## METHODS

### Study site and subjects

Field work was conducted during 98 days in total from September 2016 to April 2017. The study site is located within the *Otoch Ma’ax Yetel Kooh* reserve, close to the Punta Laguna village in the Yucatan Peninsula, Mexico (20 ° 38’ N, 87 ° 38’ W). Further details about the study site can be found in Ramos-Fernandez et al. [20]. One observer (MBJ) collected all the data on a single and habituated group of free-ranging black-handed spider monkeys (*Ateles geoffroyi*), composed of 40 individuals (see Table S1 in Supplementary material). Individuals were identified based on facial and genitalia markings and on pelage coloration. In this study we considered individuals as adults if they were at least five years old [49,50].

### Behavioural observations

Spider monkeys live in groups that constantly vary in size, cohesion and composition [37]. We considered all individuals within 30 m from one another at a given time as members of the same subgroup [30]. Observations were conducted on all individuals, older than one year old, which included 20 matures (6 males, 14 females) and 16 immatures (7 males, 9 females). Immatures were on average 3.0±1.3 years old throughout the study period. Individuals less than one year old were not included in the analysis since they were never recorded emitting the contact call types included in this study. Different subgroups were followed during consecutive days for periods of 4 to 8 hrs. per day (i.e., 32 hours per week) for a total of 548 hours. Because the size and the composition of these subgroups can be widely variable (subgroup size 8.1±5.0 individuals), the observed subgroup was semi-randomly selected, trying to equalize the total number of observation hours per individual (83.7±15 hours per individual, see supplementary material). To do so, the observer followed different subgroups on consecutive days and switched subgroups whose composition did not change after three hours in a given day. Therefore, the time spent with a given subgroup varied (3hrs16min±1hr38min).

All contact call emissions were recorded using the *all occurrence sampling* method, collecting all utterances given by any individual in the subgroup during a given time period [51]. Spider monkeys’ whinnies (including the two subtypes: low- and high-pitched whinnies [52] are easily recognizable by ear as they show a stereotyped acoustic structure composed of a series of repeated frequency modulated elements [34,53]. The identities and time of all individuals joining (fusion) and leaving (fission) the sampled subgroup were monitored continuously, allowing for the calculation of the time that each individual spent in the same subgroup with each other. To ensure the reliability of the identification of all sampled individuals, the observer (MBJ, systematically positioned in the center of the area occupied by the subgroup) was assisted by one to three experienced research assistants who were distributed all around in order to have visual access to all subgroup members and helped with the tracking and the identification of the individuals and their whinnies. We thus recorded a total of 3340 of whinnies from mature individuals and 743 whinnies from immature individuals, failing to identify callers in our observed subgroup on 157 occasions.

### Data processing

To evaluate if the changes in audience size and composition affected individual call rates, all observations were distributed into different time periods during which the composition of the studied subgroup remained stable. For example, if during an observation lasting 3 hours in total, a subgroup member left after 1 hour, two stable periods of time (one lasting 1 hour and a second one lasting 2 hours) were distinguished. For each such period of time, we counted the number of contact calls (per unit of observation time) emitted by each individual and listed the following associated characteristics of the audience:

- Subgroup size: The total number of individuals present in the audience, that is the subgroup size of the caller minus 1;
- Subgroup composition: *“Alone”* (i.e. single adult subject or mother with immature offspring; this was named *“only-mother”* when the study subject was an immature offspring knowing that immature individuals were never observed alone and never observed alone with mature males), *“same-sex”* (i.e. only mature individuals of the same sex as the caller are present), *“opposite-sex”* (i.e. only mature individuals of the opposite sex as the caller are present), *“both-sexes”* (i.e. mixed-sex audience).
- Presence or absence of mature male offspring *“with and without son”* (for mothers only).

To evaluate if the fission and fusion events affected the occurrence of contact call rates, we used the subgroup call rate as opposed to the individual call rate in order to have a sufficient time of observation under different conditions. We counted the number of calls emitted by all subgroup members 45 min before and 45min after a fission-fusion event (only fusion, only fission or both fission and fusion). We noted the “composition of the audience” (only females, only males, both sexes, alone) before and after this fission-fusion event, and we noted if the social composition remained in the same “audience sex composition” (one sex-one sex, both sexes-both sexes) or if it changed (one sex-both sexes, both sexes-one sex). For comparative purposes, we considered 90 min as an optimal period of time to see the impact of an audience change, since that is the average time that a subgroup remains stable (i.e. without fissions or fusions; [54]). All calls collected outside of these periods with fission-fusion events were considered to occur during stable periods. We included periods shorter than 90 min when for any reason we could not follow the subgroup. Consequently, when during the 45 min after the social change there was any other social change, the time of the fission-fusion event was prolonged, we noted the total duration.

### Statistical analysis

To test the effects of size and sex composition of audience on individual contact call rate, we built a generalized linear mixed model (GLMM) per age-sex category, conducting a total of four analyses for different individual categories: mature females, mature males, immature females and immature males. For each model, fixed effects were the subgroup composition categories and the number of individuals. To assess the effects of presence (1) or absence (0) of a mature son, we ran an additional GLMM. Caller identities and the date of observation were included as random factors for all models. Since our dependent variable was a rate, we ran GLMMs using a Poisson distribution with a log-link function [55]. For all the GLMM models, we compared a null model including only the random factors with a full model including all fixed effects and their interactions. We compared the null and full models using a likelihood ratio test (LRT) with the ANOVA function [56]. We selected the full model as the final if the LRT was significant and only then performed Tukey posthoc comparisons. We used the *lme4, visreg* and *multcomp* packages[57,58] in R (R Core Team 2016).

To test whether call rate varied with fission-fusion events (where periods were categorized into fusion only, fission only, both fission and fusion and stable) and whether call rate varied with changes in “audience sex composition” (one sex-one sex, one sex-both sexes, both sexes-one sexe, both sexes-both sexes), we conducted two general linear models (GLM) tests. In addition, we explored whether call rate varied before and after fission-fusion events or changes in “audience sex composition”. We ran a linear mixed model (LMM) with fission-fusion event, changes in audience sex composition and time (before and after social events or changes in audience sex composition) as fixed effects, call rate as dependent variable, and identity of the event as random effect. We included in the model all main effects and the interactions between “fission-fusion event” and the “before or after the event” (45min before and after change) and “changes in audience sex composition” and “before or after the event”. Because call rate residuals did not meet the assumptions of no significant outliers and normal distribution, we log-transformed call rate to meet both assumptions. The variances were homogeneous (Levene’s test: p = 0.1 and 0.9, respectively). We report partial eta squared (η2) and Tukey post hoc tests for ANOVA tests. We used IBM SPSS Statistics (v.25) to run the ANOVAs and LMM.

The complete data set and the full statistical code have been deposited in the Figshare data repository at the following address: https://figshare.com/s/e97dc3930ce827dd103c.

All field observations were in accordance with the ethical standards and legal requirements of the National Commission for Protected Areas in Mexico. Protocols were approved by the “Dirección General de Vida Silvestre” (SEMARNAT, permit #SGPA/DGVS/1405/15).

## RESULTS

### 1.1. Effects of size and composition of audience on individual call rate

Audience size and audience composition influenced signalers differently depending on their sex and age (Table S2, Figures 1 and 2).

**Figure 1.**
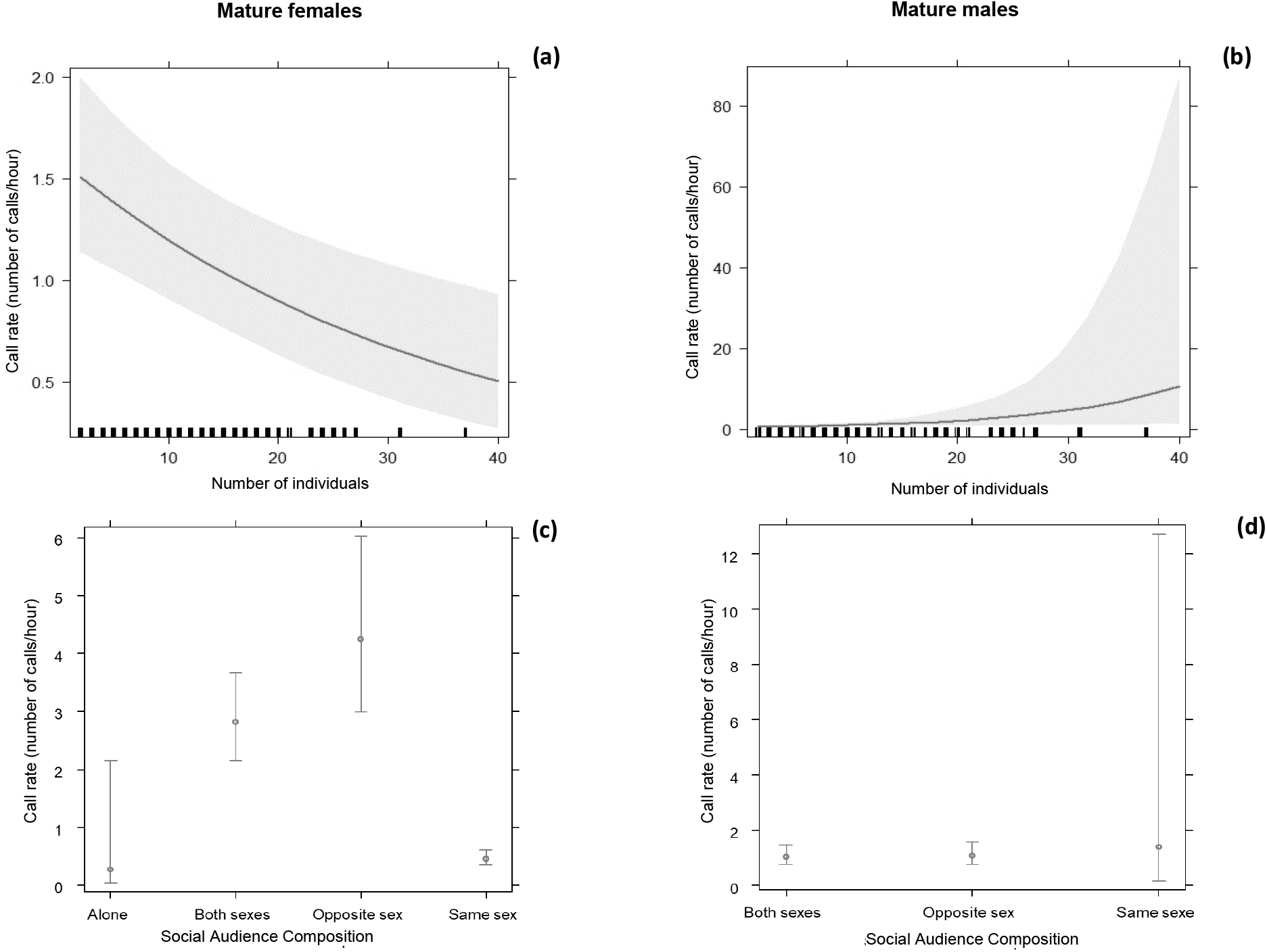
At the top of the figure, we find the effects plot that shows the expected influence of audience size on the individual call rates of (a) mature females and (b) males. The lines represent predicted means derived from the generalized linear mixed model. The gray area represents the 95% confidence interval and the rug plot at the bottom of the graphs shows the location of the audience size values. At the bottom of the figure, we find the effects plot that shows the expected influence of social audience individual call rates of (c) mature females and (d) males calling. The points are the fitted values of each category of social audience and their standard errors which are based on the covariance matrix of the estimated regression coefficients.

**Figure 2.**
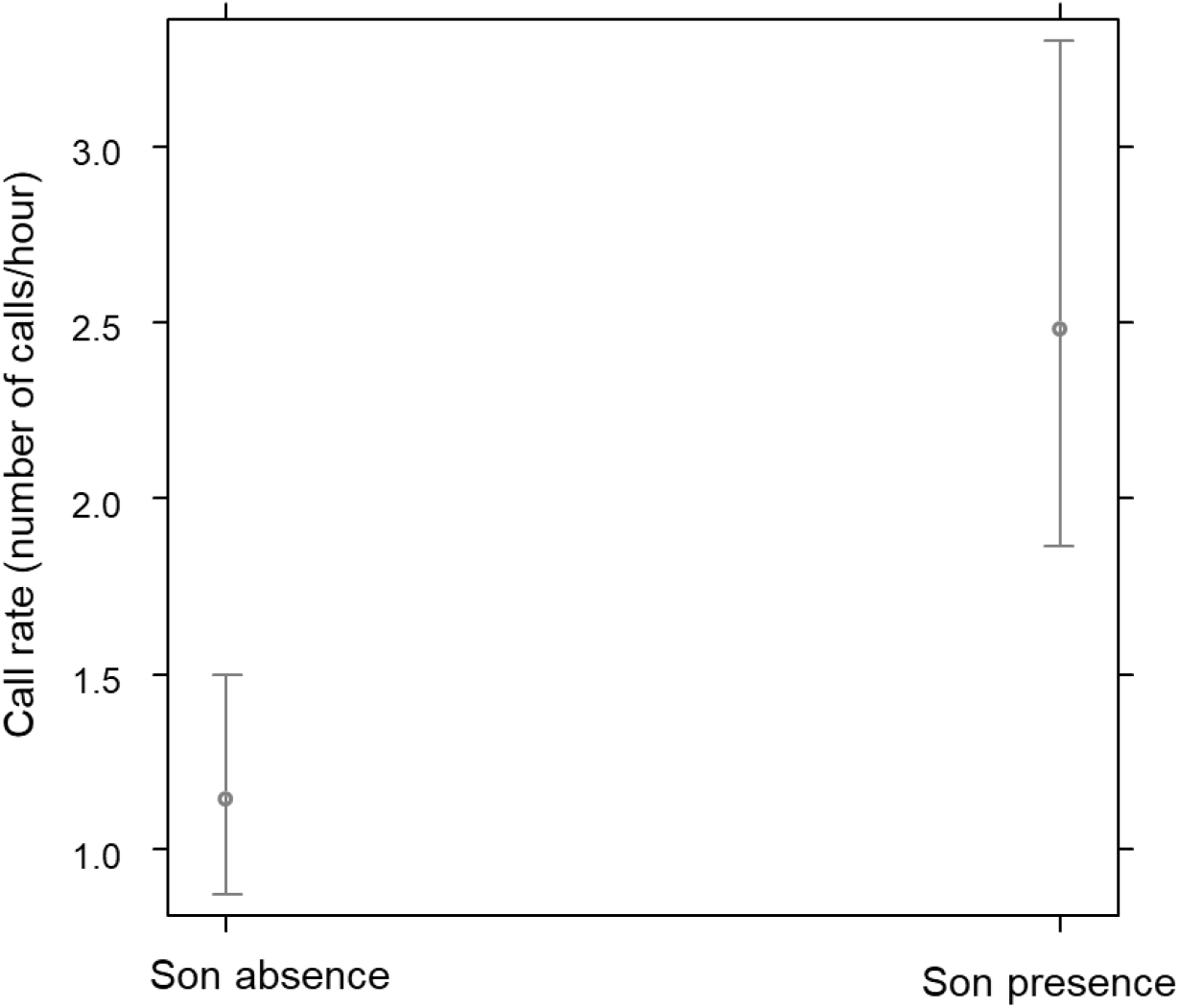
Effects plot shows the expected influence of mature son presence or absence on the call rate by mature female spider monkeys. The points are the fitted values of each category (presence or absence) and their standard errors which are based on the covariance matrix of the estimated regression coefficients.

#### Mature females

Concerning mature females, the full model was significantly different from the null model (LRT: χ^2^=1071.4; *P*<0.0001), showing that mature females’ calling rate was affected negatively by the number of individuals in the audience: the smaller the audience, the higher their call rate χ^2^=-7.62; *P*=0.005). Their calling rate was also affected by the sexual composition of the audience χ^2^=1128.1; *P*<0.0001), regardless of the audience size χ^2^=6.04; *P*=0.109). Post-hoc Tukey comparisons showed that mature females emitted calls at higher rates when only males were present in the audience, than when both males and females were present, and still lower call rates when they were with other females, their offspring or alone.

#### Mature males

Concerning mature males, the full model was also significantly different from the null model (LRT: χ2=122.81; *P*<0.0001). But conversely to mature females, mature males’ contact call rates were not affected by the size of the audience χ^2^=1.85; *P*=0.17). Similarly, to mature females, the mature males’ call rates were affected by the sexual composition of the audience: they called at higher rates when the opposite sex was present χ^2^=60.38; *P*<0.0001), regardless of the audience size χ^2^=5.25; *P*=0.07). Post-hoc Tukey comparisons showed that mature males called at higher rates when mature females were present (with or without other males) than when being in an uni-sex subgroup.

#### Immature females

Contrary to mature females and males, a sex difference was not found in immature signalers. The full models were significantly different from the null models (LRT: χ2=49.91; *P*<0.0001) showing that immature females’ χ^2^=-11.89; *P*=0.0006) similarly to mature females, were affected negatively by the size of the audience: the smaller the audience, the higher the call rates. Aditionally, immature female call rates were affected by the sexual composition of the audience but in a direction opposite to adults χ^2^=19.88; *P*=0.0002) with a significant interaction between social composition of the audience and audience size χ^2^=49.91; *P*=0.04). Immature females presented higher call rates when they were with their mother or with other members of the same sex than when in the presence of males.

#### Immature males

In the same way as immature females, the full model was significantly different from the null models (LRT: χ2=302.22; *P*<0.0001), showing that immature males’ call rates χ^2^=15.42; *P*<0.0001) were affected negatively by the size of the audience: the smaller the audience, the higher the call rates. Similar to immature females call rates were affected by the sexual composition of the audience in a direction opposite to adults χ^2^=166.79; *P*<0.0001) regardless of the audience size χ^2^=4.05; *P*=0.26). In the same way, immature male call rates called more when they were with their mother or with other members of the same sex than when being in presence of other females.

### 2. The effects of a specific individual’s presence on contact call rates

The presence of a specific individual within the audience also influences mature females’ contact call rates. The full model was significantly different from the null model (LRT: χ2=105.11; *P*<0.0001), showing that mature females’ call rates were significantly affected by the presence of their mature son in the audience χ^2^=10.59; *P*<0.0001), calling at higher rates when their mature male offspring were present in the subgroup than when they were not (Table S2, Fig. 2).

### 3. Effects of fission-fusion events on subgroup call rate

#### Call rate during fission-fusion, fission, fusion or no events

At the subgroup level, call rates are higher than at the individual level (Fig. 3) as they represent the sum of individual calling rates, which range as in Fig. 1 and Fig. 2. These rates varied significantly depending on the occurrence of fission-fusion events (F_3, 165_ = 33.6, p < 0.0001, η2 = 0.38). The post hoc test showed that subgroup call rate was significantly higher during periods when both fissions and fusions occurred than during periods when only fusion or fissions occurred. We observed the lowest call rates during fission events, which were not statistically different from periods with no fissions or fusions (Table S3, Fig. 3).

**Figure 3.**
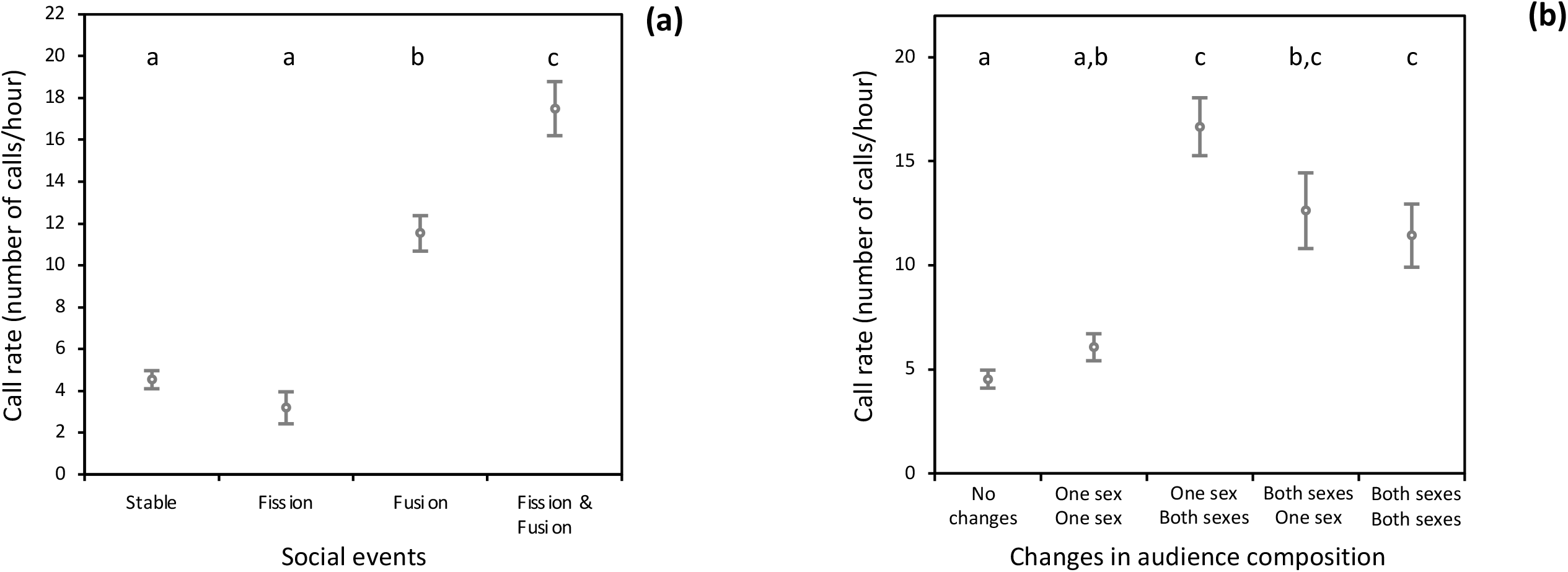
Plots showing the effects of social events and changes in audience composition on call rate. Error bars show the standard error. Letters indicate the results of the post hoc test, where groups with the same letters indicate nonsignificant differences.

#### Call rate depending on the change of subgroup composition

We also found that call rate varied significantly with changes in audience composition during fission and fusion events (F_4, 164_ = 22.3, p < 0.0001, η2 = 0.88). The post hoc test showed that call rate was higher when there was a change from one sex to two sexes and when both sexes remained in the group, intermediate when there were no changes in sex audience and for changes from two sexes to one sex, and lower when there were no audience changes in the group (Table S4, Fig. 3).

The LMM showed that call rate varied with fission-fusion events and changes in audience sex composition (F_2,135_ = 32.4, p < 0.0001; and F_3,135_ = 6.6, p < 0.0001, respectively), the effect of the before or after the event variable and its interactions with other factors were not significant (Time: F_1,135_ = 1.1, p = 0.07; fisison-fusion event*before or after the event: F_2,135_ = 1.5, p = 0.2; and changes in audience sex composition* before or after the event: F_3,135_ = 1.4 p = 0.2), suggesting that call rate did not vary before and after fission-fusion events and that the effects of type of fission-fusion event and changes in audience sex composition are similar regardless of whether one looks at the 45 minutes before or after the event.

## DISCUSSION

We studied functional and mechanistic aspects of vocal flexibility in spider monkeys, a species with high levels of social complexity according to several metrics [20,43,59]. We evaluated whether individual spider monkeys vary their emission of contact calls with the size and composition of the subgroup they are in, and whether calling at the subgroup level increases during periods where fission and fusion events occur. We found that audience of subgroup size and composition significantly affected the contact call rates of all age-sex classes but, as predicted, in an age- and sex-dependent way. Our first hypothesis was partially validated: call rates of mature females and immature individuals of both sexes decreased when the subgroup size increased. This was not the case for the mature males’ call rate. In support of our second hypothesis, we found that mature females and males increased their call rates when there was at least one individual of the opposite sex in the subgroup. Immature individuals were also affected by the sexual composition of the audience, but the pattern was the opposite as the adults’, with more calls when they were with their mother or with other members of the same sex. In support of our third hypothesis, we found that mature females called more when their mature sons were present in the audience. Finally, in concordance with our fourth hypothesis, contact call rates were higher during periods with higher social instability (during fission-fusion events) than during more stable periods, especially when there was a change in sex composition.

We predicted that large audiences would inhibit females from calling but stimulate males, mainly because females are affected more than males by food competition, and they are less invested in managing their social relationships [48]. Adult females indeed showed lower call rates when the number of individuals in the audience increased. Adult females seem to be more involved than males in repelling potential competitors and avoiding aggressive interactions [60]. Given that the intensity of competition is impacted by group size and that large groups go through more tense situations [61], a possible interpretation is that females are actually using whinnies to regulate subgroup size depending on food availability. This was actually one of the first suggested functions of the whinny [62] but the evidence so far has been equivocal [30,52]. A simpler explanation, based on the cohesion function of whinnies, is that females are less prone to losing touch with the rest of the subgroup when in larger subgroups, simply because more individuals are easier to keep track of by the noise they make while moving or by their own whinnies. Smaller subgroups may require higher rates of calling by individuals simply to stay in touch with the rest of the subgroup.

As expected, contact call rates of immature individuals presented the same pattern as their mothers. This goes in line with previous findings showing that immature spider monkeys’ associations with other group members are primarily determined by their mother’s [47]. However, it is also possible that immatures call more in small groups for different reasons than their mother; for example, that the mother’s calls motivated their offspring to vocalize. Because adult males are known to be more socially active than adult females [45,50], we predicted that larger subgroups would trigger more calling from them. But we did not find this. It is possible that adult males are less sensitive than adult females and immatures to the size composition of the audience because their social interactions with other group members are generally relaxed and face lower levels of feeding competition. Further studies should investigate whether the presence of individuals with more or less preferred social bonds or kinship influence male calling.

We also found that contact call rates were influenced by the sexual composition of the audience. Interestingly, while adult males and females called more in the presence of the opposite sex, immatures called more in the presence of a same-sex partner. This supports a function of contact calls in the regulation of encounters and spacing mediation between the sexes. Our results are different to those obtained by Dubreuil et al [32], who showed that adult female spider monkeys called more in larger subgroups. However, they also found that mature females (the study had insufficient calls for male analysis) were more likely to vocalize in the first place when the number of males increased, then when the number of females increased, and lastly when the number of males decreased or fissioned her subgroup, but not when the number of females decreased. Spider monkeys tend to form sex-segregated subgroups [40,45] and encounters between sexes (during subgroup fusions) can be tense, with both sexes prone to receiving aggression from other adults within their group [63]. Increasing the rates of whinnies in presence of the opposite sex may reduce the risk of aggressive interactions by facilitating the location of others and spacing out appropriately [32]. In chimpanzees, the presence of certain males was the most important parameter in the regulation of female call rates [64]. Intra-sex competition could be another factor responsible for the tension observed during inter-sexes reunions, and individuals could call to attract allies or identify themselves to avoid aggression. Competition between males for access to females is attested by high rates of appeasing embracing behaviors among males during sex encounters [39]. Competition between females is also observed with new immigrant females being more vulnerable to attack from resident females [46].

The fact that the immatures’ pattern differs from that of mature individuals could be due to the different social interest from males and females but also the interest they may have in individuals of the same sex as a reference to socialize. For example, immature males presented more affiliative interactions to adult males than do immature females [47]. Vocal usage certainly follows social development, as immatures typically show same-sex preferential bonding before developing heterosexual relationships [65]. In spider monkeys, strong male–male bonds begin early in life [80].

As predicted, mature females presented higher contact call rates when mature sons were present in the audience. The presence of an important social partner is a strong influence on an individual’s decision to produce calls [66]. More calls in the presence of kin partners are frequently found in primates [67,68]. Our findings support the idea that, in spider monkeys, female-male close relationships in adulthood are kin-biased [47]. Cases where females provide care and protect mature sons, and vice versa, can be found in .the spider monkey literature [47,69,70]. The presence of a mature son may also help dissuade aggression directed toward the mother herself or toward younger offspring and therefore, be a factor in promoting a mother’s overall reproductive success [47]. Also supporting these results, adult females may be more prone to respond to the whinnies of their juvenile offspring than any other juveniles [71]. This suggests that contact calls in spider monkeys also play a role in mother-offspring spatial coordination.

We have analyzed contact calling rates at the individual and subgroup levels. Individual calling rates differed between contexts, albeit by small amounts. These small differences could certainly have a functional impact if every recipient has a certain probability of behaving in a certain way in response (approaching, retreating, calling in response) to every vocalization it hears. In the long term, and over all individuals that could be recipients, there would be an effect in terms of the fission-fusion dynamics and the maintenance of social relationships between callers and recipients. The fact that females call more when in smaller subgroups, and those males call equally regardless of subgroup size, could imply that subgroups of all sizes would maintain an approximately constant call rate. However, when we analyzed the effect of fission and fusion events on this subgroup calling rate, we identified sources of variation that may be related to cohesion and spacing functions of whinnies, in the context of a dynamic subgroup composition. Indeed, social instability affected contact call rates at this subgroup level. We registered more calling in subgroups during periods with fission and fusion events, when social instability was higher, than during events when subgroups only fissioned or during periods without changes. This result is consistent with previous studies suggesting that contact calls allow individuals to obtain information about subgroup members’ locations [30], and can serve a spacing function [32] regulating fission-fusion events. During periods with high uncertainty, a flexible use of whinnies, which contain information about individual identity, might be helping to identify and locate individuals who, depending on their social relationship with the caller, might approach or retreat from it. In our study, subgroups called at higher rates when individuals may have been trying to obtain information from the other individuals that were joining the subgroup (e.g. identity or location), that is during a fusion. Additionally, more calls were registered when subgroups changed their sex composition (from one to two sexes, or vice versa) than when sex composition remain stable. It seems, then, that there is a more evident “negotiation” based on whinnies during periods of greater tension and/or uncertainty, such as the arrival of individuals of the opposite sex.

We did not find significant differences between subgroup call rates emitted 45 min before and 45 min after fission-fusions, probably due to the fact that the change in subgroup composition is a continuous process, rather than an instantaneous event. Therefore, we cannot conclude that the emission of calls before or after fusion or fission could influence the individuals to join or leave from the subgroup. Complementary studies would be necessary, taking into account the position of two subgroups before they fuse and the exact time and distance between all individuals.

In terms of the possible mechanisms underlying vocal flexibility, our results on audience effects support the idea of voluntary use of vocalizations in non-human primates. The decision of an individual to vocalize or remain silent could be shaped across learned contingencies and depends on the constant evaluation of current circumstances, comprising the relationship quality between caller and receivers, the caller’s present motivational state, and their particular socio-ecological interests [2]. Individuals could decide which of many contextual cues are relevant, and that may require elaborate cognitive processes. For example, female calling behavior in chimpanzees, who also have high degrees of fission-fusion dynamics and a marked difference in the social status of males and females [72], was moderated by social inhibition, which increased with the number of bystanders [64]. Inhibitory skills are associated with social complexity as they prevent inappropriate responses in dynamic social environments[59,73]. Vocal inhibition is also found in chimpanzees when females do not produce copulation calls while other females are nearby [74], and in capuchins who do not emit food calls when the risk of competition is high [25].

Spider monkeys have been shown to perform at comparable levels to chimpanzees in inhibitory control laboratory tasks [59], leading to the suggestion that comparable degrees of fission-fusion dynamics in both taxa require similar inhibitory control of behaviors that might be inappropriate in particular social settings within a widely varying social environment [59]. Here we have found that, despite their limited vocal repertoire, spider monkeys appear to be skilled at modifying call usage in different social contexts. They could also be varying other aspects of their calls depending on the social context. For example, there is evidence that they are able to modify their calls as a function of the distance between callers and recipients, as individuals emitted and respond with low frequency whinnies when callers are outside a subgroup (separated by long distances) to facilitate vocal contact [33].

From a functional perspective, our findings suggest that: (1) contact call usage in monkeys is flexible and context-dependent, (2) contact calls are signals that may serve several non-mutually exclusive functions and the composition of subgroups could largely be regulated by whinnies during fission and fusion events in the context of sex-segregated groups. Individuals are able to adapt to the nuances of the immediate and changing social scene characteristic of a society with a high degree of fission–fusion dynamics [75] and (3) contact calls are used to reduce the uncertainty about subgroup composition and serve a socio-spatial cohesion function. From a mechanistic perspective, our results add to the growing number of empirical demonstrations of audience effects in animals [76]. Further research is needed to determine whether the presence of specific individuals (other kin individuals, or close affiliates) can also influence call rates [77,78]. Our results support the idea that studying the potential effects of audience in species with complex social systems could allow us to predict which specific social variables may be responsible for selection on new or more complex signals and their adaptive functions [2].

## Acknowledgments

We are grateful to R.I.M. Dunbar and two anonymous reviewers for their comments and suggestions on a previous version of the manuscript. We thank the Comisión Nacional de Áreas Naturales Protegidas of Mexico (CONANP) and the Environmental agency of Mexico (Dirección General de Vida Silvestre, SEMARNAT) for the permission we received to work at the Otoch Ma’ax Yetel Kooh reserve. We are very grateful to the field assistants Augusto, Eulogio and Macedonio Canul. We thank Filippo Aureli, Colleen M. Schaffner and Laura G. Vick for sharing the management of the long-term project at Punta Laguna.

## Authors’ contributions

All authors contributed to the conceptualization, funding acquisition, project administration, formal analysis, writing original draft. Briseño-Jaramillo also performed the data curation.

## ANNEX 1

**Table S1.**
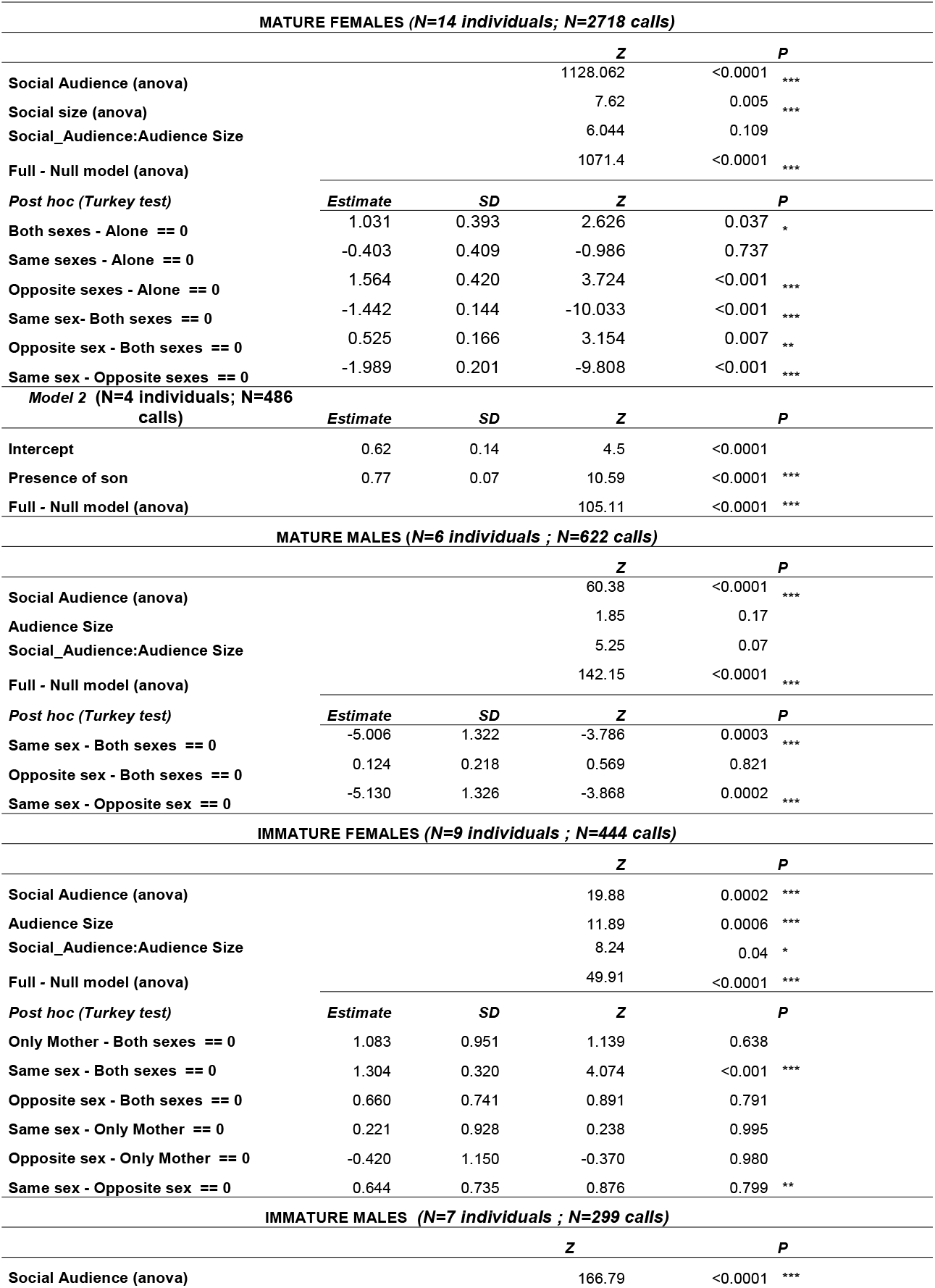
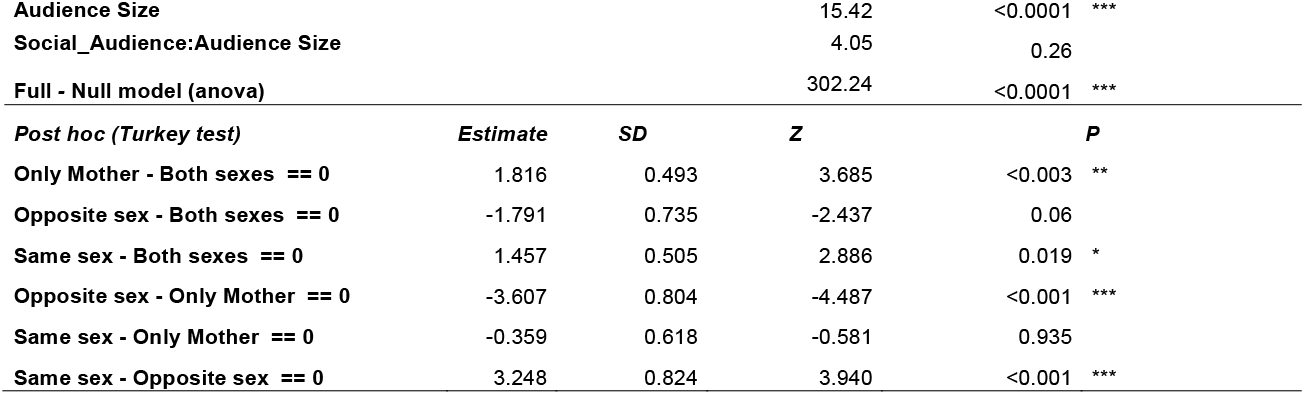
Results of each statistical model and Post hoc (Turkey test) comparisons. All assumptions for the GLMM analyses were met (collinearity and overdispersion). Results of each statistical model and Post hoc (Turkey test) comparisons. All assumptions for the GLMM and GLM analyses were met (collinearity and overdispersion).

**Table S2.**
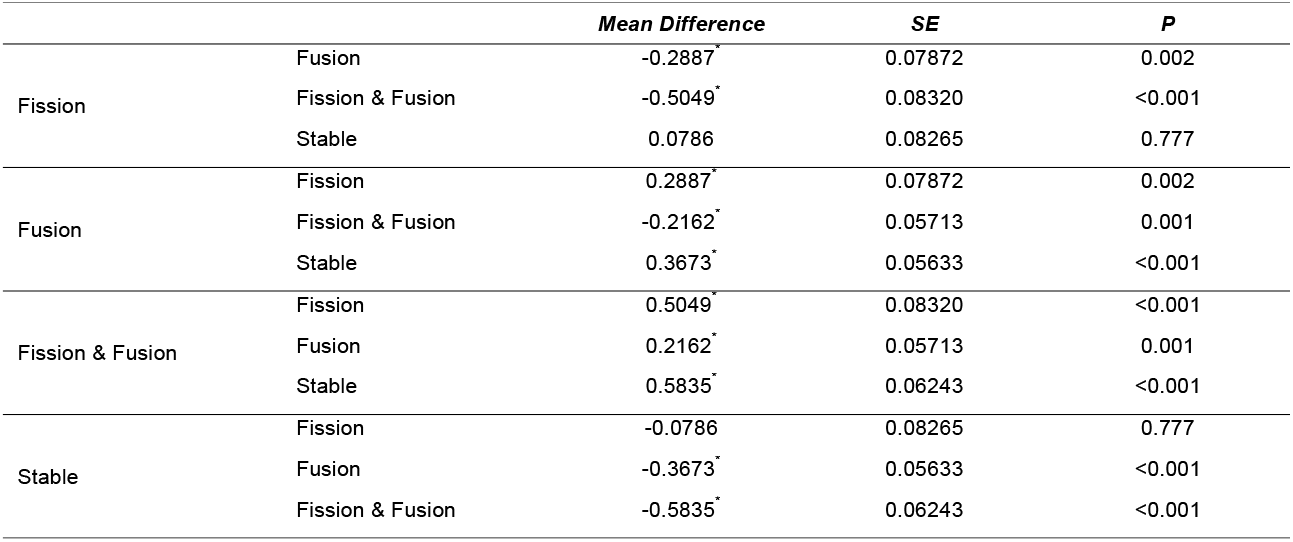
Post-hoc comparison for the analysis of social events vs. call rate.

**Table S3.**
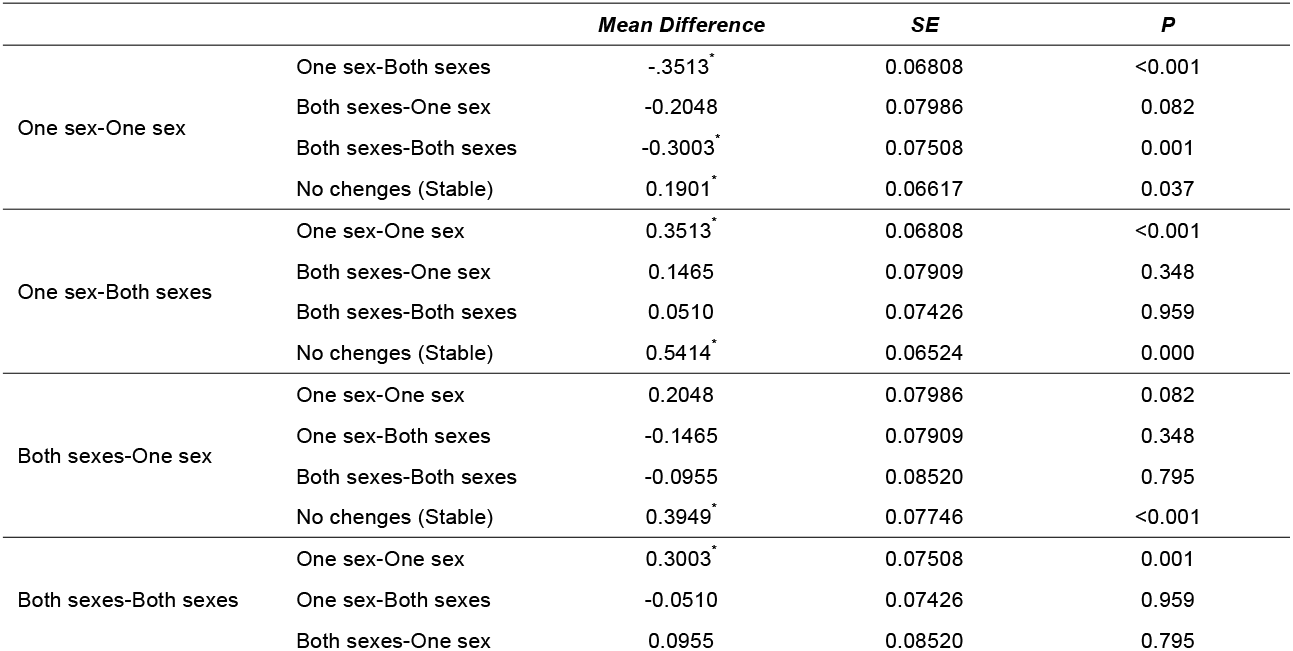
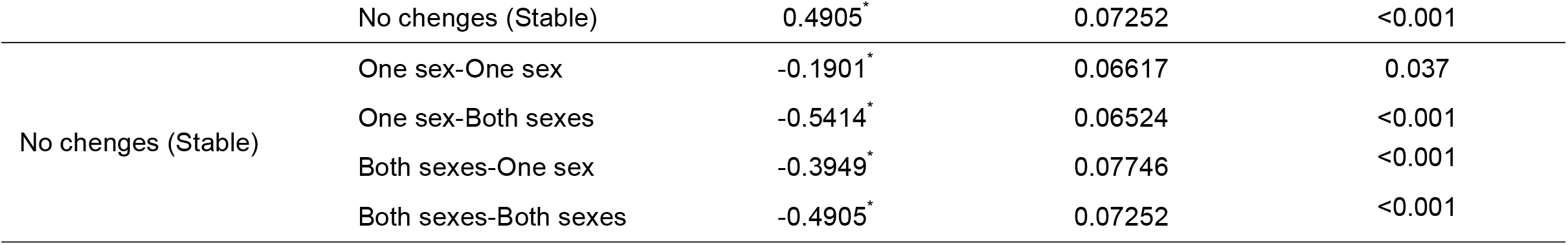
Post-hoc comparison for the analysis of changes in audience composition on call vs. call rate.

