## Supplementary material Tables S1, S2, S3 for "Flexible use of contact calls in a species with high fission-fusion dynamics"

**Table S1.** Group composition and individual characteristics of spider monkeys (*Ateles geoffroyi*) in the Otoch Ma’ax Yetel Kooh reserve, Yucatan Peninsula, Mexico.

| **Individual Code (identity)** | **Sex^a^** | **Age^b^** | **Total observation (hrs)** |
| --- | --- | --- | --- |
| FL | F | MA | 73.8 |
| VE | F | MA | 103.6 |
| CH | F | MA | 106.5 |
| LO | F | MA | 100.8 |
| KL | F | MA | 59.5 |
| JA | F | MA | 89.5 |
| VA | F | MA | 93.3 |
| HI | F | MA | 84.7 |
| PC | F | MA | 81.6 |
| TG | F | MA | 76.7 |
| ML | F | MA | 87.6 |
| AE | F | MA | 71.6 |
| EL | F | MA | 65.2 |
| MI | F | MA | 58.4 |
| TK | F | IM | 96.5 |
| LE | F | IM | 100.8 |
| LB | F | IM | 59.3 |
| PU | F | IM | 86.2 |
| TZ | F | IM | 82.1 |
| ES | F | IM | 100.8 |
| FR | F | IM | 84.7 |
| XT | F | IM | 59.1 |
| PN | F | IM | 76.7 |
| TU | M | MA | 87.9 |
| JN | M | MA | 88.4 |
| EG | M | MA | 92.4 |
| MS | M | MA | 80.5 |
| KO | M | MA | 64.5 |
| WB | M | MA | 59.1 |
| DG | M | IM | 87.7 |
| SH | M | IM | 82.9 |
| VK | M | IM | 106.3 |
| DL | M | IM | 73.8 |
| AS | M | IM | 90.5 |
| AP | M | IM | 71.6 |
| NA | M | IM | 106.5 |

^a^ F = Female, M = Male

^b^ MA = Mature; IM = Immature

**Table S2.** Results of each statistical model and Post hoc (Turkey test) comparisons.

All assumptions for the GLMM analyses were met (collinearity and overdispersion).

| **MATURE FEMALES *(N=14 individuals; N=2718 calls)*** | | | | | |
| --- | --- | --- | --- | --- | --- |
|  |  |  | ***Z*** | ***P*** | |
| **Social Audience (anova)** |  |  | 1128.062 | <0.0001 | *** |
| **Social size (anova)** |  |  | 7.62 | 0.005 | *** |
| **Social_Audience:Audience Size** |  |  | 6.044 | 0.109 |  |
| **Full *-* Null model (anova)** |  |  | 1071.4 | <0.0001 | *** |
| ***Post hoc (Turkey test)*** | ***Estimate*** | ***SD*** | ***Z*** | ***P*** | |
| **Both sexes - Alone == 0** | 1.031 | 0.393 | 2.626 | 0.037 | * |
| **Same sexes - Alone == 0** | -0.403 | 0.409 | -0.986 | 0.737 |  |
| **Opposite sexes - Alone == 0** | 1.564 | 0.420 | 3.724 | <0.001 | *** |
| **Same sex- Both sexes == 0** | -1.442 | 0.144 | -10.033 | <0.001 | *** |
| **Opposite sex - Both sexes == 0** | 0.525 | 0.166 | 3.154 | 0.007 | ** |
| **Same sex - Opposite sexes == 0** | -1.989 | 0.201 | -9.808 | <0.001 | *** |
| ***Model 2*  (N=4 individuals; N=486 calls)** | ***Estimate*** | ***SD*** | ***Z*** | ***P*** | |
| **Intercept** | 0.62 | 0.14 | 4.5 | <0.0001 |  |
| **Presence of son** | 0.77 | 0.07 | 10.59 | <0.0001 | *** |
| **Full *-* Null model (anova)** |  |  | 105.11 | <0.0001 | *** |
| **MATURE MALES (*N=6 individuals ; N=622 calls)*** | | | | | |
|  |  |  | ***Z*** | ***P*** | |
| **Social Audience (anova)** |  |  | 60.38 | <0.0001 | *** |
| **Audience Size** |  |  | 1.85 | 0.17 |  |
| **Social_Audience:Audience Size** |  |  | 5.25 | 0.07 |  |
| **Full *-* Null model (anova)** |  |  | 142.15 | <0.0001 | *** |
| ***Post hoc (Turkey test)*** | ***Estimate*** | ***SD*** | ***Z*** | ***P*** | |
| **Same sex - Both sexes == 0** | -5.006 | 1.322 | -3.786 | 0.0003 | *** |
| **Opposite sex - Both sexes == 0** | 0.124 | 0.218 | 0.569 | 0.821 |  |
| **Same sex - Opposite sex == 0** | -5.130 | 1.326 | -3.868 | 0.0002 | *** |
| **IMMATURE FEMALES *(N=9 individuals ; N=444 calls)*** | | | | | |
|  |  |  | ***Z*** | ***P*** | |
| **Social Audience (anova)** |  |  | 19.88 | 0.0002 | *** |
| **Audience Size** |  |  | 11.89 | 0.0006 | *** |
| **Social_Audience:Audience Size** |  |  | 8.24 | 0.04 | * |
| **Full *-* Null model (anova)** |  |  | 49.91 | <0.0001 | *** |
| ***Post hoc (Turkey test)*** | ***Estimate*** | ***SD*** | ***Z*** | ***P*** | |
| **Only Mother - Both sexes == 0** | 1.083 | 0.951 | 1.139 | 0.638 |  |
| **Same sex - Both sexes == 0** | 1.304 | 0.320 | 4.074 | <0.001 | *** |
| **Opposite sex - Both sexes == 0** | 0.660 | 0.741 | 0.891 | 0.791 |  |
| **Same sex - Only Mother == 0** | 0.221 | 0.928 | 0.238 | 0.995 |  |
| **Opposite sex - Only Mother == 0** | -0.420 | 1.150 | -0.370 | 0.980 |  |
| **Same sex - Opposite sex == 0** | 0.644 | 0.735 | 0.876 | 0.799 | ** |
| **IMMATURE MALES  *(N=7 individuals ; N=299 calls)*** | | | | | |
|  |  |  | ***Z*** | ***P*** | |
| **Social Audience (anova)** |  |  | 166.79 | <0.0001 | *** |
| **Audience Size** |  |  | 15.42 | <0.0001 | *** |
| **Social_Audience:Audience Size** |  |  | 4.05 | 0.26 |  |
| **Full *-* Null model (anova)** |  |  | 302.24 | <0.0001 | *** |
| ***Post hoc (Turkey test)*** | ***Estimate*** | ***SD*** | ***Z*** | ***P*** | |
| **Only Mother - Both sexes == 0** | 1.816 | 0.493 | 3.685 | <0.003 | ** |
| **Opposite sex - Both sexes == 0** | -1.791 | 0.735 | -2.437 | 0.06 |  |
| **Same sex - Both sexes == 0** | 1.457 | 0.505 | 2.886 | 0.019 | * |
| **Opposite sex - Only Mother == 0** | -3.607 | 0.804 | -4.487 | <0.001 | *** |
| **Same sex - Only Mother == 0** | -0.359 | 0.618 | -0.581 | 0.935 |  |
| **Same sex - Opposite sex == 0** | 3.248 | 0.824 | 3.940 | <0.001 | *** |

**Table S3.** Post-hoc comparison for the analysis of fission-fusion events vs. call rate. Results of each statistical model and Post hoc (Turkey test) comparisons. All assumptions for the GLMM and GLM analyses were met (collinearity and overdispersion).

|  |  | ***Mean Difference*** | ***SE*** | ***P*** |
| --- | --- | --- | --- | --- |
| Fission | Fusion | -0.2887^*^ | 0.07872 | 0.002 |
|  | Fission & Fusion | -0.5049^*^ | 0.08320 | <0.001 |
|  | Stable | 0.0786 | 0.08265 | 0.777 |
| Fusion | Fission | 0.2887^*^ | 0.07872 | 0.002 |
|  | Fission & Fusion | -0.2162^*^ | 0.05713 | 0.001 |
|  | Stable | 0.3673^*^ | 0.05633 | <0.001 |
| Fission & Fusion | Fission | 0.5049^*^ | 0.08320 | <0.001 |
|  | Fusion | 0.2162^*^ | 0.05713 | 0.001 |
|  | Stable | 0.5835^*^ | 0.06243 | <0.001 |
| Stable | Fission | -0.0786 | 0.08265 | 0.777 |
|  | Fusion | -0.3673^*^ | 0.05633 | <0.001 |
|  | Fission & Fusion | -0.5835^*^ | 0.06243 | <0.001 |

**Table S4.** Post-hoc comparison for the analysis of changes in audience composition on call vs. call rate. Results of each statistical model and Post hoc (Turkey test) comparisons.

All assumptions for the GLMM and GLM analyses were met (collinearity and overdispersion).

|  |  | ***Mean Difference*** | ***SE*** | ***P*** |
| --- | --- | --- | --- | --- |
| One sex-One sex | One sex-Both sexes | -.3513^*^ | 0.06808 | <0.001 |
|  | Both sexes-One sex | -0.2048 | 0.07986 | 0.082 |
|  | Both sexes-Both sexes | -0.3003^*^ | 0.07508 | 0.001 |
|  | No chenges (Stable) | 0.1901^*^ | 0.06617 | 0.037 |
| One sex-Both sexes | One sex-One sex | 0.3513^*^ | 0.06808 | <0.001 |
|  | Both sexes-One sex | 0.1465 | 0.07909 | 0.348 |
|  | Both sexes-Both sexes | 0.0510 | 0.07426 | 0.959 |
|  | No chenges (Stable) | 0.5414^*^ | 0.06524 | 0.000 |
| Both sexes-One sex | One sex-One sex | 0.2048 | 0.07986 | 0.082 |
|  | One sex-Both sexes | -0.1465 | 0.07909 | 0.348 |
|  | Both sexes-Both sexes | -0.0955 | 0.08520 | 0.795 |
|  | No chenges (Stable) | 0.3949^*^ | 0.07746 | <0.001 |
| Both sexes-Both sexes | One sex-One sex | 0.3003^*^ | 0.07508 | 0.001 |
|  | One sex-Both sexes | -0.0510 | 0.07426 | 0.959 |
|  | Both sexes-One sex | 0.0955 | 0.08520 | 0.795 |
|  | No chenges (Stable) | 0.4905^*^ | 0.07252 | <0.001 |
| No chenges (Stable) | One sex-One sex | -0.1901^*^ | 0.06617 | 0.037 |
|  | One sex-Both sexes | -0.5414^*^ | 0.06524 | <0.001 |
|  | Both sexes-One sex | -0.3949^*^ | 0.07746 | <0.001 |
|  | Both sexes-Both sexes | -0.4905^*^ | 0.07252 | <0.001 |
